# Genomic characterization reveals high clonal redundancy in two *Acropora cervicornis* nurseries in the Dominican Republic

**DOI:** 10.64898/2026.09.11.750210

**Authors:** Shamwari Anseeuw Carrasco, Kasey H. Walsh, Rebecca Garcia-Camps, Ainhoa L. Zubillaga, Aldo Croquer, Debashish Bhattacharya

## Abstract

*Acropora cervicornis* is often propagated through fragmentation (i.e., clonal propagation), a common practice implemented by restoration projects. Known as asexual propagation, this strategy may rapidly increase coral cover but reduces genetic variation. We used 2b-RAD sequencing to characterize multilocus genotypic variation among 45 *A. cervicornis* colonies maintained in two in-situ nurseries (Cap Cana, Acuario) in Punta Cana, Dominican Republic. After reference-based SNP discovery and filtering, 2,515 high-quality SNPs were retained in the dataset. Principal coordinate analysis (PCoA), hierarchical clustering, identity-by-state (IBS) distances, and relatedness estimates identified only three multilocus genets among the 45 colonies (AC1, AC2, and AC3). Of these, AC2 was the most abundant (n = 23; 51.1%), followed by AC3 (n = 14; 31.1%) and lastly AC1 (n = 8; 17.8%). Within-genet IBS distances (0.153-0.227) did not overlap with between-genet distances (0.406-0.497). AC1 and AC2 were detected in both nurseries, whereas AC3 was limited to Acuario. These results reveal substantial clonal redundancy among sampled nursery colonies that determines local reef conservation efforts. These data highlight the value of incorporating genotype identification in nursery-based coral restoration projects.

## Introduction

The Caribbean reef-building coral *Acropora cervicornis* has undergone severe population declines throughout its historical range due to disease (e.g., White Band Disease, WBD), bleaching events, hurricanes, and other disturbances (Aronson & Precht, 2001; Carpenter et al., 2008). As a result, coral restoration has become an important strategy to promote the recovery of this species, with in-situ nurseries used widely to maintain colonies for subsequent outplanting (Young et al., 2012; Lirman & Schopmeyer, 2016; Hodges & Hallock, 2025).

Coral fragmentation allows the fast-growing *A. cervicornis* to be rapidly propagated, but multiple nursery colonies may represent ramets (fragments) of the same genet rather than genetically distinct individuals (genets) (Tunnicliffe, 1981; Highsmith, 1982). In such cases, coral abundance may not reflect genotypic diversity expected in target wildlife populations. To address this issue, genetic characterization of coral nurseries is needed to identify clonal redundancy, quantify genet representation, and determine whether genetic lineages are distributed among multiple facilities. The final goal of a coral nursery is to assist recovery by providing a source of propagules for outplanting (Boström-Einarsson et al. 2020) while keeping genetic diversity of the wildlife coral populations (Shearer et al. 2009; Baums et al. 2022).

Here, we used 2b-RAD sequencing to characterize 45 *A. cervicornis* colonies maintained in two in-situ nurseries (Cap Cana, Acuario) in Punta Cana, Dominican Republic (DR). We quantified the number and relative abundance of multilocus genets, evaluated genomic support for genet assignments, and determined their distribution between nursery facilities. This study provides a genomic baseline of the genetic inventory represented among sampled colonies in these two important DR nurseries.

## Methods

### Study sites and sample collection

Sampling was conducted in May 2025 at two in-situ *A. cervicornis* nurseries in Punta Cana, DR. Forty-five tagged colonies were sampled: 19 from Cap Cana (18.475418° N, 68.388340° W) and 26 from Acuario (18.539046° N, 68.348283° W). The Acuario nursery was established approximately 15 years ago with an estimated 13 genotypes originating from several locations spanning hundreds of kilometers along the northern and southern coasts of the DR, including Sosúa, Bayahibe, Cap Cana, and Punta Rucia (pers. comm.). The Cap Cana nursery was established in September 2020 by Fundación Cap Cana using coral fragments collected from a single nearby site, but records of the number of genotypes initially introduced were unavailable (pers. comm.). The two nurseries are separated by approximately 8 km and have an average water depth of 5 m. Colonies are maintained on rope structures, domes, and arches. Approximately 2 cm of branch tissue was collected from each colony and preserved in DNA/RNA Shield (Zymo Research), under CITES Permit DO-02381, dated 22 May 2025.

### DNA extraction and 2b-RAD sequencing

Genomic DNA was extracted using the Zymo Research Quick-DNA Miniprep Kit following the manufacturer’s protocol. 2b-RAD library preparation and sequencing were performed by the vendor CD Genomics. Approximately 50ng of genomic DNA per sample was digested with *Bsa*XI, ligated to sample-specific adapters, amplified using Phusion High-Fidelity DNA Polymerase, purified by polyacrylamide gel extraction, and barcoded. Libraries were pooled and sequenced on an Illumina NovaSeq X Plus platform using 150bp paired-end sequencing. Technical replicate libraries prepared from the same DNA extractions were not available.

### Read processing and SNP discovery

Initial sequence processing was performed by CD Genomics. Paired-end reads were merged using PEAR v0.9.6 (Zhang et al., 2014), adapters were removed, and the terminal three bases were excluded to minimize ligation-site artifacts. Reads lacking the expected restriction site, containing >8% ambiguous bases, or having >80% of bases below Q30 were removed. Additional quality and length trimming was performed with Cutadapt v5.2 (Martin, 2011).

Quality-filtered reads were aligned to the *A. cervicornis* reference genome jaAcrCerv1.1 (GCA_964034985.1) using Bowtie2 v2.5.4 (Langmead & Salzberg, 2012), and alignments were sorted and indexed using SAMtools v1.22.1 (Li et al., 2009). Genotype likelihoods were estimated in ANGSD using the SAMtools model, retaining uniquely mapped reads and excluding reads flagged as bad. Bases with quality scores <25 and reads with mapping quality scores <30 were excluded. Candidate SNPs were identified using a polymorphism-test significance threshold of *P*< 1 x 10^-5^. SNPs were retained when represented in at least 34 of 45 individuals, with minor allele frequency (MAF) ≥ 0.05, and after strand-bias and heterozygote-bias filtering. Triallelic sites were excluded. The final dataset contained 2,515 SNPs. Full processing parameters and the final ANGSD command are provided in the Supplementary Methods.

### Genet identification and genetic differentiation

Pairwise identity-by-state (IBS) estimates generated in ANGSD were used to construct an IBS distance matrix. Genetic structure was evaluated in R v4.4.2 (R Core Team, 2024) using principal coordinates analysis (PCoA) and average-linkage hierarchical clustering. Multilocus genet assignments were defined from a three-cluster solution, and concordance between hierarchical clustering and the groups observed in the PCoA was evaluated. Assignment stability was assessed across dendrogram cut heights (**Fig. S1**), and pairwise IBS distances were summarized within and between inferred genets.

Pairwise relatedness (rab) was estimated using ngsRelate v2.0 (Hanghøj et al., 2019) from ANGSD-derived genotype likelihoods. Because the presence of many genetically identical ramets can influence allele-frequency estimation, the analysis was repeated using one representative ramet per genet. Genotype likelihoods and allele frequencies were re-estimated for the three representatives using the 2,515 SNP positions retained in the full analysis, and pairwise relatedness was recalculated with ngsRelate.

### Genotypic diversity and nursery distribution

The number and frequency of multilocus genets were summarized. Genotypic richness was calculated as *R* = (*G*−1) / (*N*−1), where *G* is the number of multilocus genets detected and *N* is the number of sampled ramets. Genotypic evenness was quantified using Simpson evenness *V* following Arnaud-Haond et al. (2007). The full evenness calculation is provided in the Supplementary Methods.

### Data and statistical analyses

Downstream analyses and visualization were performed in RStudio (R Core Team, 2026). Raw sequencing data was deposited in the NCBI Sequence Read Archive under BioProject PRJNA1514087, and analysis scripts and processed data necessary to reproduce the analyses are archived at https://github.com/shamwarianseeuw/DR_ACER_PopGen_2025.

## Results

### Sequencing and SNP discovery

Sequencing generated 447.3 million raw reads from the 45 colonies, of which 419.0 million (93.68%) were retained following quality filtering. Of 7,930 candidate SNPs, 2,515 (31.7%) passed all filtering criteria and were retained for downstream analyses (**Table S1**). Mean depth across the 2,515 retained SNP sites was 75.50x per sample (SD = 13.11x; range = 57.86–114.07x; **Table S2**).

### Genet identification and differentiation

Population genomic analyses identified three multilocus genets: AC1, AC2, and AC3 in the dataset (**Fig. 1**). PCoA of pairwise IBS distances showed three discrete groups corresponding to these genets (**Fig. 1a**). Average-linkage hierarchical clustering recovered the same three groups, with cluster membership agreeing with the genet assignments observed in the PCoA (**Fig. 1b**). The three-cluster solution was stable across dendrogram cut heights from 0.222 to 0.426 (**Fig. S1**).

**Fig. 1.**
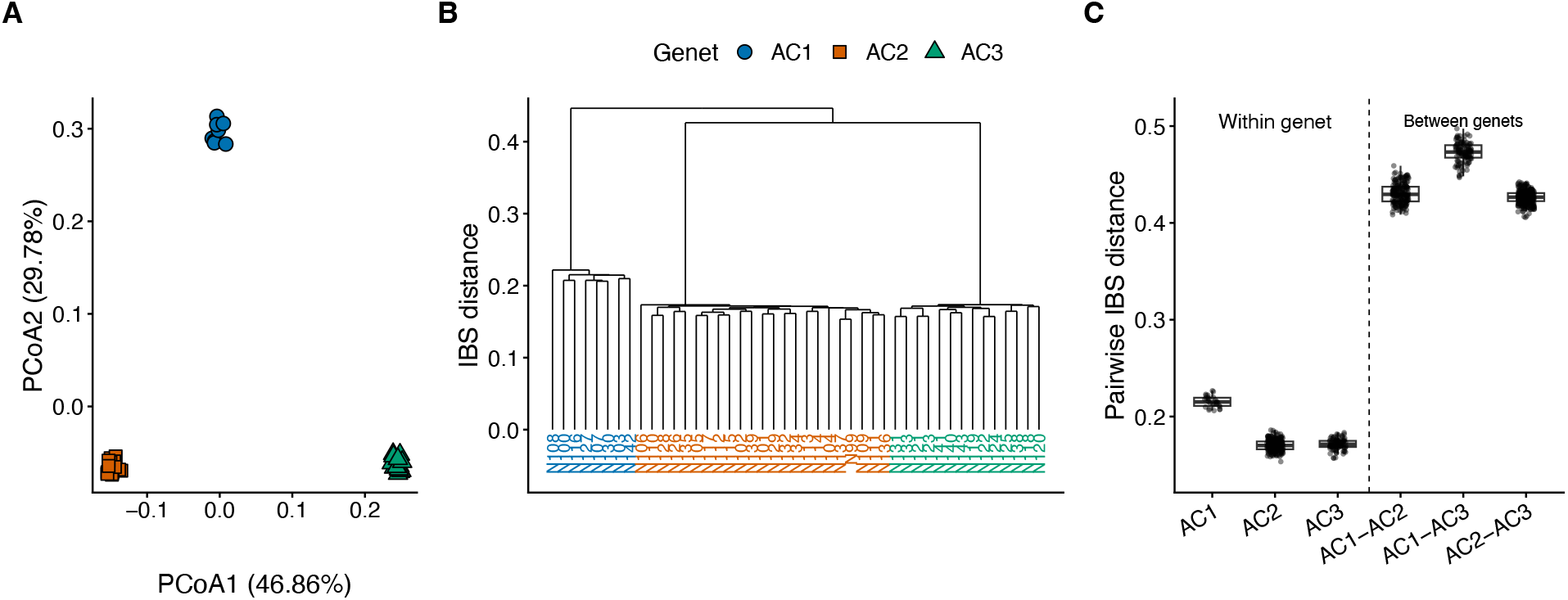
Genetic differentiation and multilocus genet assignment among 45 *Acropora cervicornis* colonies sampled from two in-situ nurseries in Punta Cana, DR. **A)** Principal coordinates analysis (PCoA) of pairwise identity-by-state (IBS) distances. PCoA1 and PCoA2 explain 46.86% and 29.78% of the variation, respectively; **B)** Average-linkage hierarchical clustering of pairwise IBS distances, recovering the same three genets (AC1, n = 8; AC2, n = 23; AC3, n = 14); and **C)** Distribution of pairwise IBS distances within and between genets.

Pairwise IBS distances showed complete separation between within and between-genet comparisons (**Fig. 1c**; **Table S3**). Within-genet distances ranged from 0.153 to 0.227, whereas between-genet distances ranged from 0.406 to 0.497. Mean within-genet distances were 0.215 for AC1, 0.170 for AC2, and 0.171 for AC3; mean between-genet distances ranged from 0.426 to 0.473.

Relatedness estimates provided additional support for these three assignments (**Table S4**). Mean within-genet r_ab_ was 0.9993 for AC1, 0.9989 for AC2, and 0.9996 for AC3, whereas between-genet estimates were effectively zero (0-3 x 10^-6^). After reducing the dataset to one representative ramet per genet and re-estimating allele frequencies, r_ab_ remained effectively zero for all three pairwise genet comparisons.

### Genotypic diversity and nursery distribution

The 45 colonies represented three genets: AC1 (n = 8; 17.8%), AC2 (n = 23; 51.1%), and AC3 (n = 14; 31.1%) (**Table 1**; **Fig. 2**). Overall genotypic richness was R = 0.045, whereas genotypic evenness was V = 0.903. Genet composition differed between nurseries. Cap Cana had two detected genets: AC1 (n = 5) and AC2 (n = 14), whereas Acuario contained all three: AC1 (n = 3), AC2 (n = 9), and AC3 (n = 14).

**Table 1.** Genotypic composition and diversity of *Acropora cervicornis* colonies sampled from the Cap Cana and Acuario in-situ nurseries in Punta Cana, DR. Values for AC1-AC3 show the number and percentage of sampled ramets assigned to each multilocus genet. *N* is the number of sampled ramets, *G* the number of detected genets, *R*, is the genotypic richness, and *V* is Simpson’s genotypic evenness (Arnaud-Haond et al., 2007).

| Nursery | $N$ | AC1, $n$ (%) | AC2, $n$ (%) | AC3, $n$ (%) | $G$ | $R$ | $V$ |
| --- | --- | --- | --- | --- | --- | --- | --- |
| Cap Cana | 19 | 5 (26.3) | 14 (73.7) | 0 (0.0) | 2 | 0.056 | 0.720 |
| Acuario | 26 | 3 (11.5) | 9 (34.6) | 14 (53.8) | 3 | 0.080 | 0.828 |
| **Overall** | **45** | **8 (17.8)** | **23 (51.1)** | **14 (31.1)** | **3** | **0.045** | **0.903** |

**Fig. 2.**
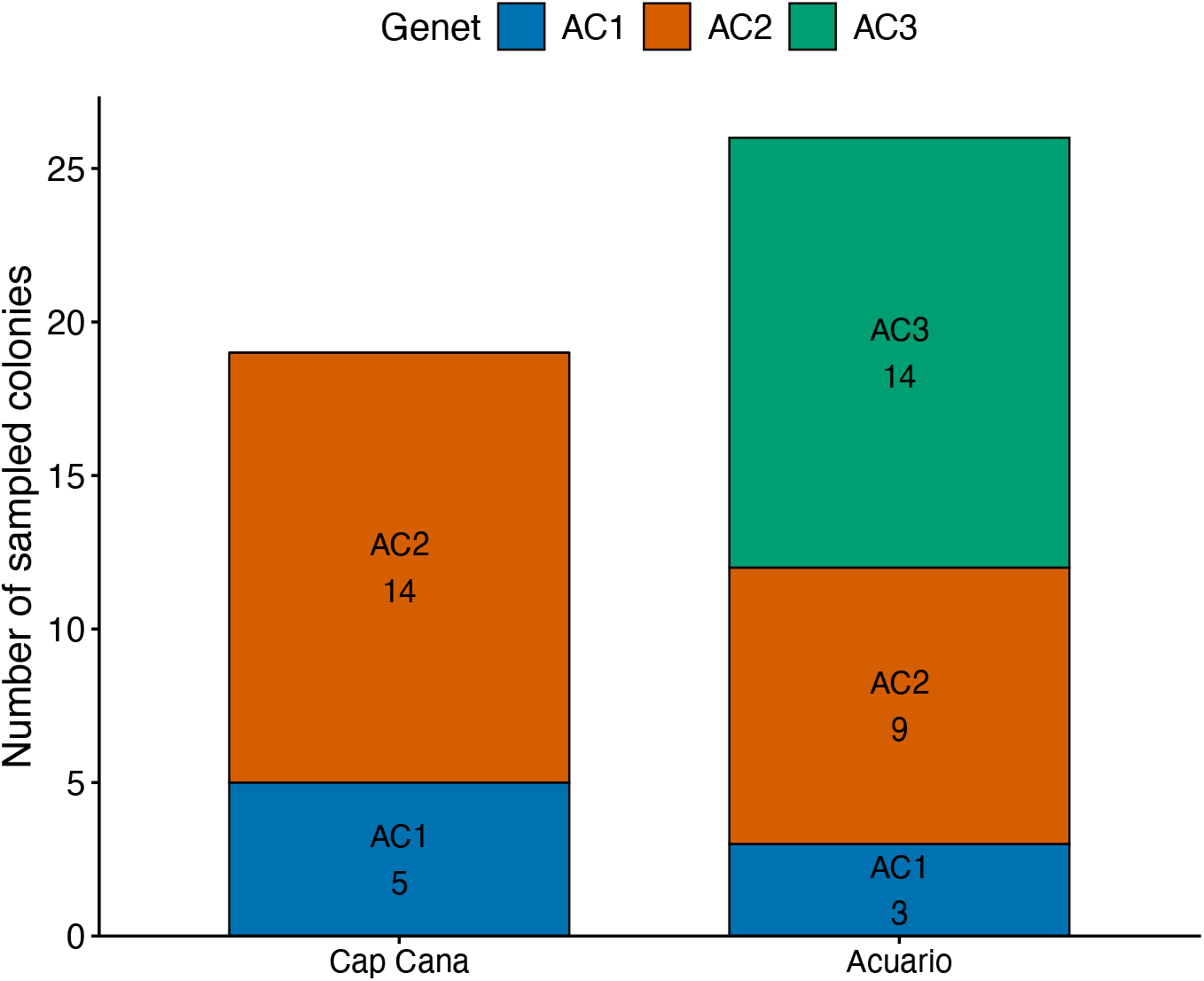
Distribution of multilocus genets among two *Acropora cervicornis* in-situ nurseries in Punta Cana, DR. The stacked bars show the number of sampled ramets assigned to each multilocus genet (AC1-AC3) within the Cap Cana and Acuario nurseries. AC1 and AC2 were detected in both nurseries, whereas AC3 was detected only at Acuario.

## Discussion

Genomic characterization of 45 *A. cervicornis* colonies from two Punta Cana nurseries identified three multilocus genets which showed a large clonal redundancy among the sampled sites. Multiple analyses independently supported the same assignments: PCoA and hierarchical clustering recovered three discrete groups, assignments were stable across a range of clustering thresholds, and within and between-genet IBS distances were non-overlapping. Pairwise relatedness neared the value of one within genets and was effectively zero among genets. These results held up when relatedness among genets was re-estimated using one representative ramet per genet. Together, these results provide strong support for AC1, AC2, and AC3 being distinct, clonally propagated genets in the two in-situ nurseries.

The recovery of three genets among 45 colonies shows the clear difference between coral colony abundance and genetic diversity in clonally propagated (i.e. fragmentated) coral nurseries. Overall genotypic richness was low (R = 0.045), although the three detected genets were relatively evenly represented (V = 0.903). For comparison, wild *A. cervicornis* populations in the U.S. Virgin Islands had substantially higher genotypic richness (R = 0.62; Nylander-Asplin et al., 2021) than observed in the sampled nursery stocks (R = 0.045), although differences in sampling and population context should be considered when comparing these values. Genet representation also differed between facilities: AC1 and AC2 were found in both nurseries, whereas AC3 was only in Acuario. The two sampled nurseries therefore did not contain the same genetic diversity, demonstrating that inventories incorporating genet identity provide useful information that cannot be captured by only relying on colony abundance values. Genetic characterization has also been used to assess the diversity of *A. cervicornis* nursery stocks elsewhere in the DR and to inform restoration planning for local practitioners (Calle-Triviño et al., 2020).

The low genotypic richness observed in this study is relevant to conservation programs in the area because maintaining genetic diversity is an important consideration when selecting and propagating *A. cervicornis* for restoration (Lirman & Schopmeyer, 2016; Calle-Triviño et al., 2020). Previous genomic studies have shown substantial genetic variation within and among surviving *A. cervicornis* populations in the Florida Reef Tract, highlighting the value of incorporating genetic data into restoration strategies (Drury et al., 2017a). Genetic identity can also impact restoration outcomes, because different *A. cervicornis* genotypes may differ in growth rate and bleaching response, recovery, and survival (Drury et al., 2017b).

Low genotypic richness was observed in the sampled nurseries despite the Acuario nursery being established with coral fragments originating from geographically separated locations across the DR. Because genotype information for the original fragments is unavailable, it is unclear what factors led to the current genet composition. One possibility is that the original fragments had low genetic diversity despite being collected from different locations. In contrast, differences in survivorship over the years, including during major bleaching events such as the 2023 Caribbean mass bleaching event (Hoegh-Guldberg et al., 2023) may have reduced the number of genets represented in the nurseries. Propagation practices and coral fragment transfers among facilities may have further influenced the current genet composition. Therefore, the three genets detected here may reflect both the genetic composition of the original nursery stock and the subsequent survival and propagation of particular geneotypes.

This study was limited to two restoration nurseries and does not characterize *A. cervicornis* genetic diversity throughout the DR. Comparisons with surrounding wild populations and other nurseries, such as the study conducted by Calle-Treviño et al. (2020) are needed to determine whether additional genets occur regionally and how representative the sampled nursery stocks are of local diversity. Technical replicate libraries were also unavailable, preventing estimation of a protocol-specific genotyping-error threshold. Nevertheless, the complete separation of within- and between-genet IBS distances, stability of genet assignments across clustering thresholds, and agreement among clustering, ordination, and relatedness analyses provide multiple lines of evidence supporting the proposed genet assignments. These data establish a genomic baseline for genetic diversity in the sampled DR nurseries and underline why genet identification is necessary to build successful coral restoration programs.

## Acknowledgments

Primary funding for this study was provided by the National Philanthropic Trust (23-7825575) in a grant awarded to Debashish Bhattacharya. This work was also supported by a grant from the USDA National Institute of Food and Agriculture Hatch Formula (NJ01180) awarded to DB. Samples were collected and exported to the US under CITES Permit DO-02381. We wish to acknowledge the help of research interns in the Dominican Republic Marine Innovation Hub for aid in processing samples, the captains of the research vessels who led the different sampling trips, and Fundación Cap Cana for allowing us to sample their nursery. This project is part of the International Climate Initiative (IKI). The Federal Ministry for the Environment, Nature Conservation and Nuclear Safety (BMU) supports this initiative based on a decision adopted by the German Bundestag.

## Data Availability

Raw sequence reads generated in this study are available through the NCBI Sequence Read Archive under BioProject PRJNA1514087. Scripts used for sequence processing, population genomic analyses, statistical analyses, and figure generation are available at https://github.com/shamwarianseeuw/DR_ACER_PopGen_2025.

