## Supplemental materials and methods for "Genomic characterization reveals high clonal redundancy in two *Acropora cervicornis* nurseries in the Dominican Republic"

**Supplementary Methods**

*Detailed sequence-processing and ANGSD parameters*

Following initial sequence processing by CD Genomics, additional quality and length trimming was performed with Cutadapt v5.2 using -q 15,15 -m 25. Quality-filtered reads were aligned to the *Acropora cervicornis* reference genome jaAcrCerv1.1 (GCA_964034985.1) using Bowtie2 v2.5.4 with the parameters --local -L 16 --score-min L,16,1 --no-unal. Alignments were sorted and indexed using SAMtools v1.22.1.

Genotype-likelihood estimation, SNP discovery, and IBS calculations were performed in ANGSD using the following parameters:

-GL 1

-uniqueOnly 1

-remove_bads 1

-minMapQ 30

-minQ 25

-dosnpstat 1

-doHWE 1

-sb_pval 1e-5

-hetbias_pval 1e-5

-skipTriallelic 1

-minInd 34

-snp_pval 1e-5

-minMaf 0.05

-doMajorMinor 1

-doMaf 1

-doCounts 1

-makeMatrix 1

-doIBS 1

-doCov 1

-doGeno 8

-doBCF 1

-doPost 1

-doGlf 2

-P 1

Only uniquely mapped reads were retained, reads flagged as bad were removed, and minimum mapping- and base-quality thresholds of MAPQ ≥ 30 and Q ≥ 25, respectively, were applied. Candidate SNPs were required to have qualifying data in at least 34 of 45 sampled colonies, MAF ≥ 0.05, and a polymorphism-test significance of P < 1 x 10^-5^. Sites failing the ANGSD strand-bias or heterozygote-bias filters (-sb_pval 1e-5 and -hetbias_pval 1e-5) were excluded, as were triallelic sites. An exploratory ANGSD quality-control analysis was performed before selection of the final filtering thresholds. Parameters from that exploratory analysis were used only to assess sequencing and coverage characteristics and were not used to generate the final SNP dataset.

*SNP filtering and site-retention criteria*

ANGSD identified 7,930 candidate SNPs before completion of final SNP-level quality control. Of these candidate sites, 3,812 satisfied the minimum-individual criterion (minInd ≥ 34), and 3,741 satisfied both minInd ≥ 34 and MAF ≥ 0.05. These counts represent the numbers of candidate SNPs satisfying the respective criteria and do not constitute sequential filtering steps. The complete filtering procedure additionally incorporated the mapping- and base-quality thresholds, SNP significance threshold, strand-bias and heterozygote-bias filters, and exclusion of triallelic sites described above. Following all filters, 2,515 of 7,930 candidate SNPs (31.7%) were retained for downstream analyses (**Table S4**).

*Genotypic richness and evenness*

Genotypic richness was calculated as described in the main Methods. For calculation of Simpson's genotypic evenness (*V*), finite-sample-corrected Simpson diversity was calculated as *D* = 1 - ∑*_i_n_i_* (*n_i_* - 1) / *N* (*N*-1), where *n_i_* is the number of ramets assigned to genet *i*. The minimum and maximum possible diversity for *N* ramets distributed among *G* genets were calculated as *D*_min_ = (*G*-1) (2*N*-*G*) / *N* (*N*-1), and *D*_max_ = *N* (*G*-1) / *G* (*N*-1). Simpson's genotypic evenness was then calculated as *V* = (*D*-*D*_min_) / (*D*_max_-*D*_min_). Values approaching 1 indicate increasingly even representation of ramets among the observed genets.

The complete computational workflow and analysis scripts are available at https://github.com/shamwarianseeuw/DR_ACER_PopGen_2025.

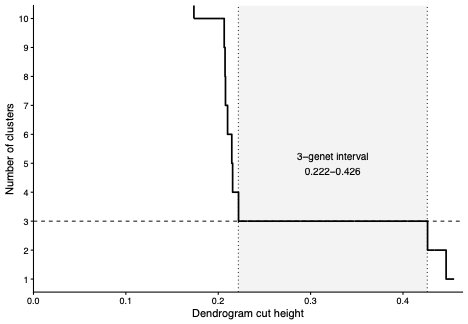

**Fig. S1** Sensitivity of multilocus genet assignment to hierarchical-clustering cut height among 45 *Acropora cervicornis* colonies sampled from two in-situ nurseries in Punta Cana, DR. The number of clusters recovered from the average-linkage dendrogram based on pairwise identity-by-state (IBS) distances is shown across dendrogram cut heights. A stable three-genet solution occurred between heights of 0.222 and 0.426. Throughout this interval, the same three groups were recovered with identical membership: AC1 (n = 8), AC2 (n = 23), and AC3 (n = 14). The final within-genet merge occurred at 0.222, whereas the subsequent merge at 0.426 reduced the solution from three to two clusters.

**Table S1.** Summary of SNP discovery and filtering in the reference-based *Acropora cervicornis* dataset. Candidate SNPs were identified in ANGSD using a polymorphism-test significance threshold of P < 1 × 10⁻⁵. Counts for the minimum-individual (minInd ≥ 34) and minor-allele-frequency (MAF ≥ 0.05) criteria show the number of candidate SNPs satisfying each criterion independently and jointly and therefore do not represent sequential filtering steps. The complete filtering pipeline additionally excluded sites failing strand-bias or heterozygote-bias thresholds and triallelic sites. After all filters, 2,515 of 7,930 candidate SNPs (31.7%) were retained for downstream analyses.

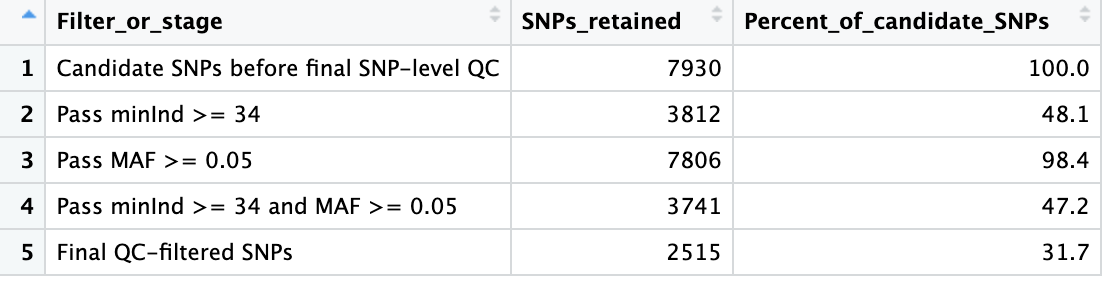

**Table S2.** Sample metadata, multilocus genet assignments, and sequencing depth for 45 *Acropora cervicornis* colonies sampled from the Cap Cana and Acuario nurseries in Punta Cana, DR. For each colony, the table reports sample identifier, nursery of origin, inferred genet (AC1-AC3), the number of the 2,515 retained SNP sites with qualifying sequence coverage, and mean sequencing depth across all 2,515 retained SNP sites. Depth was calculated from reference-aligned reads passing minimum mapping-quality (MAPQ ≥ 30) and base-quality (Q ≥ 25) thresholds; retained SNP sites without qualifying coverage in an individual were included as zero-depth observations. Across the 45 colonies, mean depth was 75.50x ± 13.11 SD (median = 73.61x; range = 57.86-114.07x).

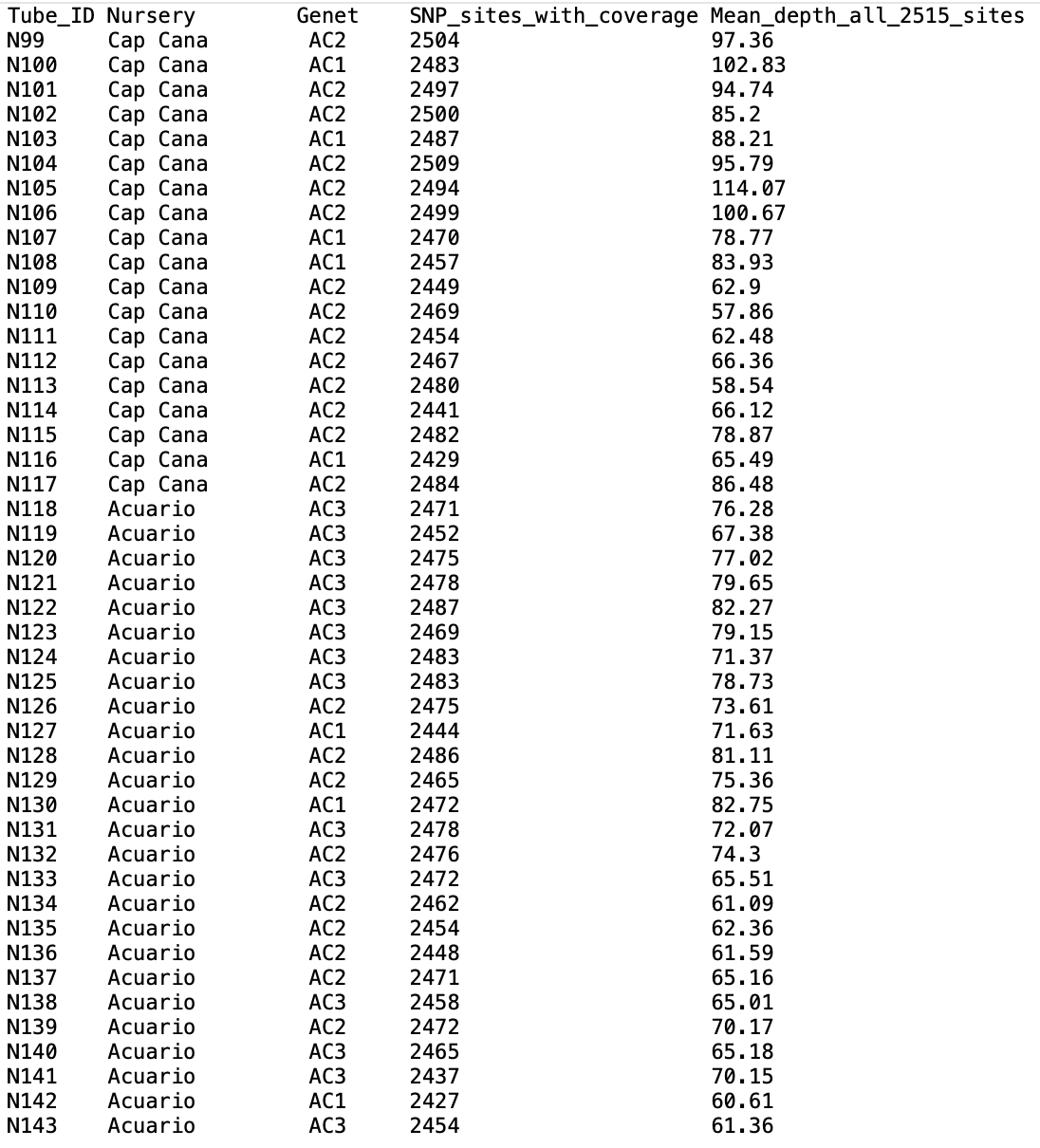

**Table S3.** Pairwise identity-by-state (IBS) distances within and between the three multilocus genets identified among 45 *Acropora cervicornis* colonies sampled from two in-situ nurseries in Punta Cana, DR. Pairwise comparisons are summarized by the number of comparisons (n), mean, standard deviation (SD), median, minimum, maximum, and first and third quartiles (Q25 and Q75). Across all 372 within-genet comparisons, IBS distances ranged from 0.1534 to 0.2266, whereas the 618 between-genet comparisons ranged from 0.4056 to 0.4973. The within- and between-genet distributions did not overlap.

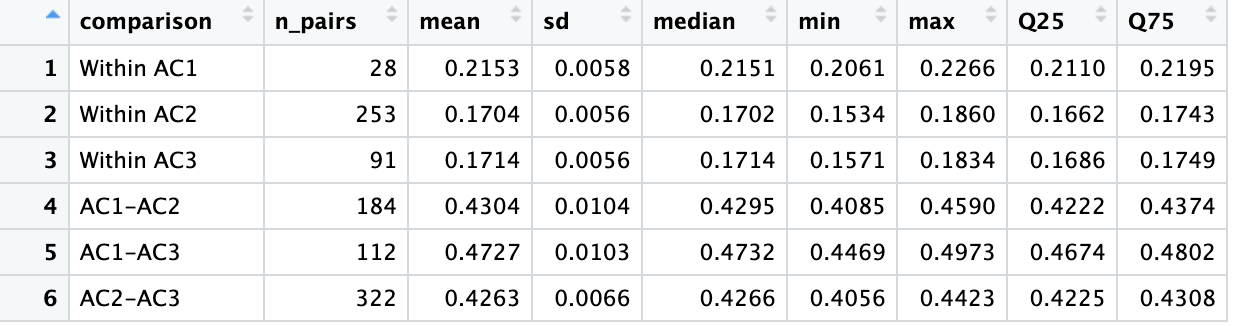

**Table S4.** Pairwise relatedness among multilocus genets identified in the *Acropora cervicornis* nurseries. **(A)** Pairwise ngsRelate relatedness estimates (r_ab_) from the full dataset of 45 colonies, summarized within and between the three inferred genets. Values are reported as the number of pairwise comparisons, mean, standard deviation, minimum, and maximum. **(B)** Pairwise relatedness estimated after reducing the dataset to one representative ramet per genet (AC1: N130; AC2: N128; AC3: N120) and re-estimating allele frequencies from the reduced dataset. The reduced analysis was used to assess whether the strong clonal representation in the full dataset influenced estimates of relatedness among genets.

**A**

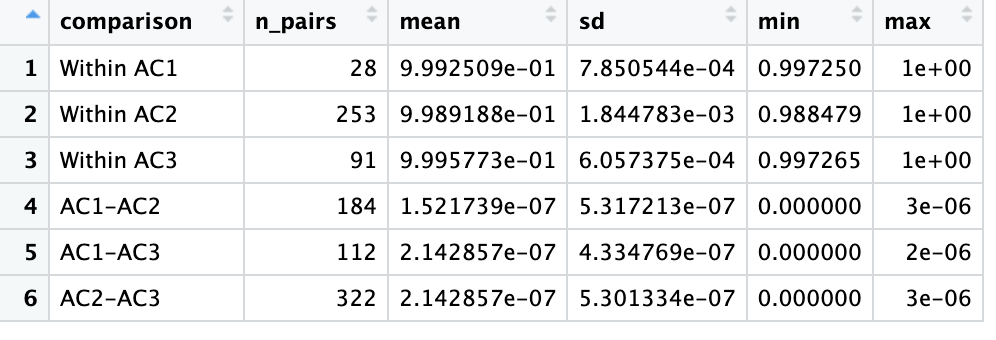

**B**

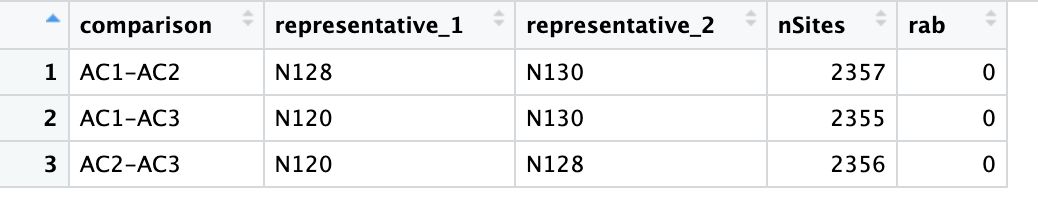
